# Cyclical compression loading is the dominant mechanoregulator of synovial joint morphogenesis

**DOI:** 10.1101/2023.02.09.527957

**Authors:** Josepha Godivier, Elizabeth A. Lawrence, Mengdi Wang, Chrissy L. Hammond, Niamh C. Nowlan

## Abstract

Mechanical stimuli arising from fetal movements are critical factors underlying joint growth. Abnormal fetal movements negatively affect joint shape features with important implications for joint function and health, but the mechanisms by which mechanical forces due to fetal movements influence joint growth are still unclear. In this research, we integrated cell-level data into a novel mechanobiological model of zebrafish jaw joint morphogenesis to identify links between the mechanical stimuli arising from movement and patterns of growth. Larval zebrafish jaw joint growth patterns were quantified from tracked cell-data at several successive developmental stages in the presence or absence of movements. Pharmacological immobilisation, prior to the onset of jaw movements, resulted in growth rate decreases which were stronger along the ventrodorsal axis. Simulations of joint morphogenesis, based on the quantified cell-level data and which integrated mechanical stimuli arising from simulated jaw movements, were used to test hypotheses relating specific mechanical stimuli with the local changes in size and shape. Different types of mechanical stimulation were incorporated into the simulation to provide the mechanoregulated component on growth in addition to the baseline (non mechanoregulated) growth which occurs in the immobilised animals. We found that the magnitude of compression experienced during joint motion when included as the stimulus for mechanoregulated growth could not predict the real, normally loaded shaped joints. However, when the dynamic changes caused by the application of cyclical compression was implemented as the stimulus for mechanoregulated growth, the sizes and shapes of joints were correctly simulated. We conclude therefore that the cyclical application of compression loading due to the dynamic nature of fetal movements underlies the mechanoregulation of prenatal joint morphogenesis. Our results provide a fundamental advance in our understanding of mechanoregulation of the developing joint and increase our understanding of the origins of conditions such as hip dysplasia and arthrogryposis.

**Author summary:** The mechanical forces caused by fetal movements are important for normal development of the skeleton, and in particular for joint shape. Several common developmental musculoskeletal conditions such as developmental dysplasia of the hip and arthrogryposis are associated with reduced or restricted fetal movements. Paediatric joint malformations impair joint function and can be debilitating. To understand the origins of such conditions, it is essential to understand how the mechanical forces arising from movements influence joint growth and shape. In this research, we used a computational model of joint growth applied to the zebrafish jaw joint to study the impact of fetal movements on joint growth and shape. We find that the cyclical application of compression loading is critical to the normal growth and shape of the developing joint. Our findings implicate that dynamic compression must be targeted when developing strategies for the treatment of musculoskeletal conditions through targeted physiotherapy.

## Introduction

Fetal movements are critical for healthy skeletal development, and abnormal movement *in utero* is associated with several conditions affecting babies in which the joint does not acquire the correct shape. Developmental dysplasia of the hip and arthrogryposis are two examples of such conditions, both of which can have lasting health consequences including the early onset of osteoarthritis [1, 2]. When skeletal muscle is absent or non-contractile in animal models, skeletal malformations include the loss of interlocking joint shape features and fusion of the skeletal elements in some (but not all) joints [3-16]. In pharmacologically paralysed chicks, for example, the femoral epiphyses are narrower both at the level of the knee [12] and of the hip [4, 12] with a loss of the acetabulum depth [4], while in muscleless-limb mice, the femoral condyles, though smaller that control littermates, are abnormally protruding [3]. Joint morphogenesis, the process by which joints acquire their shapes, is determined by co-ordinated cell activities including proliferation [5, 7, 8, 17] and changes in cell orientation, size and intercalation [5, 7, 15, 18]. The mechanisms through which mechanical loading from fetal movements regulates the cellular activities underlying joint morphogenesis are unclear. Chondrocyte proliferation [5, 7, 8, 18] and intercalation [5, 7] are impaired in the absence of embryonic movement. In paralysed zebrafish jaw joints and in muscleless-limb mice elbow joints, chondrocytes are generally smaller and rounder than those of controls and have an altered orientation, indicating cell immaturity [8, 15, 19]. The organisation of chondrocytes into columns in the growth plate, which contributes to rudiments’ elongation, is inhibited in animal models of abnormal fetal movements [7, 15]. Despite observations at the tissue and cellular level, the mechanisms by which fetal movements influence joint morphogenesis are still unclear.

Insights on cartilage mechanoregulation can be gained by studying the effects of mechanical loads on cartilage *in vivo/in ovo*, cartilage explants *ex vivo* or chondrocytes *in vitro. In vivo* [5], *in ovo* [4, 9] and *in vitro* [20] studies have shown that the development of functioning joints depends on the timing and duration of movement. While early movements, prior to joint cavitation (the physical separation of the skeletal elements), are crucial for the separation of joint elements [4, 5, 7, 9, 12, 15], short periods of immobility after cavitation has taken place have only minor influence on joint morphology [4]. However, long periods of immobilisation, even after cavitation has occurred, result in marked shape changes which can lead to joint fusion in most extreme cases in chick limbs [4, 9, 12] and larval zebrafish jaws [5, 7, 15]. Fetal chick knees cultured *in vitro* showed that the duration of loading is an important factor influencing joint growth and morphogenesis, with longer durations resulting in more normally developed joints [20]. Tissue engineering research has interrogated the effects of dynamic loading on chondrocytes *in vitro*, either through direct compression or hydrostatic pressure loading. Direct compression loading promotes extracellular matrix synthesis and tissue material properties as reviewed in [21, 22]. Significant increases in glycosaminoglycan (GAG) content were reported when dynamic compression was applied to juvenile bovine chondrocytes compared to unloaded controls [23, 24]. Increased production of collagen due to dynamic loading is less evident; a review compiling results from 63 studies reported that more studies reported no increase or a decrease in collage content when dynamic compression was applied, as opposed to unloaded controls, than studies which reported a positive effect [21]. Meanwhile, cyclic hydrostatic loading significantly increases ECM synthesis with upregulation of both GAG and collagen productions [25-27]. Compressive dynamic loading also leads to increased chondrocyte proliferation compared with free-swelling controls [28]. The duration of loading is a key parameter for the positive effect of dynamic compression on chondrogenesis, with long-duration studies tending to find increased GAG or collagen contents compared to short-duration studies as reviewed in [21]. In contrast with dynamic loading, static compression has a degenerative effect on chondrocyte metabolism leading to, for example, decreased GAG content [29-31]. While valuable insights have been gained on the specific parameters influencing chondrocyte mechanoregulation, mainly *in vitro*, the biomechanical regulation of the cells underlying joint morphogenesis remains largely unclear.

Mechanobiological simulations offer a means to integrate mechanical and biological information to bring about insights not possible with traditional approaches [32, 33]. Mechanobiological models of joint growth and morphogenesis have indicated that mechanical stimuli arising from joint motion can predict the emergence of shape features seen under normal or altered loading conditions [34-38]. For example, when simulating hip joint growth, asymmetric loading conditions resulted in shape alterations of the femoral head [34-36] and the acetabulum [36] which were characteristic of shape features seen in hip dysplasia [34, 36] or cerebral palsy [35]. Modelling muscle atrophy due to brachial plexus birth injury enabled the prediction of deformed glenohumeral joint shapes as seen in children [38]. A recent study of the regenerating axolotl humerus correlated interstitial pressure, driven by cyclic loading, with joint growth and shape changes [37]. However, previous mechanobiological models [34, 35, 37-39], including our own [36, 40], have not used accurate data for cell-level inputs. The biological contributions to morphogenesis has been assumed to be proportional to chondrocyte density which was considered either uniform across the rudiment [34, 35, 37, 38] or decreasing proportional to distance from the joint line [36, 39, 40]. A range of different biophysical stimuli (peak, minimum or average hydrostatic stress [34-36, 39-43], octahedral shear stress [34, 35, 41, 43, 44] and interstitial fluid pressure [37]) have been corelated with growth and morphogenesis, but a framework to quantitatively compare the relationships between specific stimuli and developmental change is lacking. To further explore the complex relationship between mechanical loading and joint morphogenesis, precise and specific characterisation of the contributions of cell-level dynamics to joint growth is necessary, in addition to modelling frameworks which allow the testing of hypotheses relating specific biophysical stimuli to developmental change.

Over recent years, progress has been made in characterising the cellular dynamics involved in tissue growth. Spatial morphometric analyses were conducted on light-sheet images of the embryonic murine tibia, revealing that a number of cell morphological changes and growth strategies contribute to growth plates’ expansion, especially highly spatially-dependent cell volume expansions [45]. Quantification of tissue growth based on cell lineage tracking data in the developing chick limb bud [46] and the Drosophilia wing disc [47] showed that spatially and temporally heterogeneous growth patterns coupled with growth anisotropy are major drivers of tissue morphogenesis. Recent work from our group reported that growth in the zebrafish jaw joint exhibits pronounced anisotropy likely influenced by cell orientation [48]. Integrating accurately quantified cell-level data into new mechanobiological models of joint growth will greatly deepen our understanding of the mechano-regulatory processes involved.

In this research, we aimed to identify the causal relationship between specific aspects of the biomechanical stimuli arising from embryonic movements and the patterns of joint growth and morphogenesis. We quantified zebrafish jaw joint morphogenesis, and the underlying cell activities, in the presence of normal and altered mechanical environments, and found that growth rates were diminished along a specific anatomical axis (the ventrodorsal axis) between free-to-move and immobilised larvae. Next, we designed a mechanobiological model of zebrafish jaw joint morphogenesis integrating quantified cell-activities to test hypotheses on how biomechanical stimuli arising from movements promote growth pattern changes. It emerged that, rather than the magnitude of compressions experienced during movement alone, the most likely stimulus for mechano-regulated growth is the dynamic changes arising from cyclic compression of the joint elements.

## Results

### Immobilisation leads to growth rate alterations along specific anatomical axes which reflect joint shape changes

The shapes of Meckel’s cartilage (MC) joint elements (shown in Fig 1A and B) from larvae immobilised from day 3 post fertilisation and from free-to-move larvae (controls) were measured at 4, 4.5 and 5 days post fertilisation (dpf). While there were no significant differences in shape measurements between free-to-move and immobilised groups, or between timepoints within each group, the free-to-move larvae exhibited higher increases of MC length over the whole timeframe compared to immobilised larvae as seen with average shape outlines in Fig 1Cb, d. MC length increased by approximately 37% from 4 to 5 dpf in the free-to-move larvae compared to an average increase of 15% in the immobilised larvae over the same timeframe (Fig 1D). Immobilised MC depth remained almost constant from 4 to 5 dpf whereas free-to-move MC depth markedly increased over the same timeframe (43% increase in free-to-move larvae, compared to 6% decrease in the immobilised larvae) (Fig 1Cb, d and Fig 1D). Free-to-move MC width at the level of the joint increased slightly over the investigated timeframe (11% increase) whereas the immobilised MC width remained almost unchanged over time (4% increase; Fig 1Ca, c and Fig 1D). Therefore, growth of the depth of the MC was most severely affected by the absence of jaw movements, with growth of MC length and width less affected.

**Fig 1.**
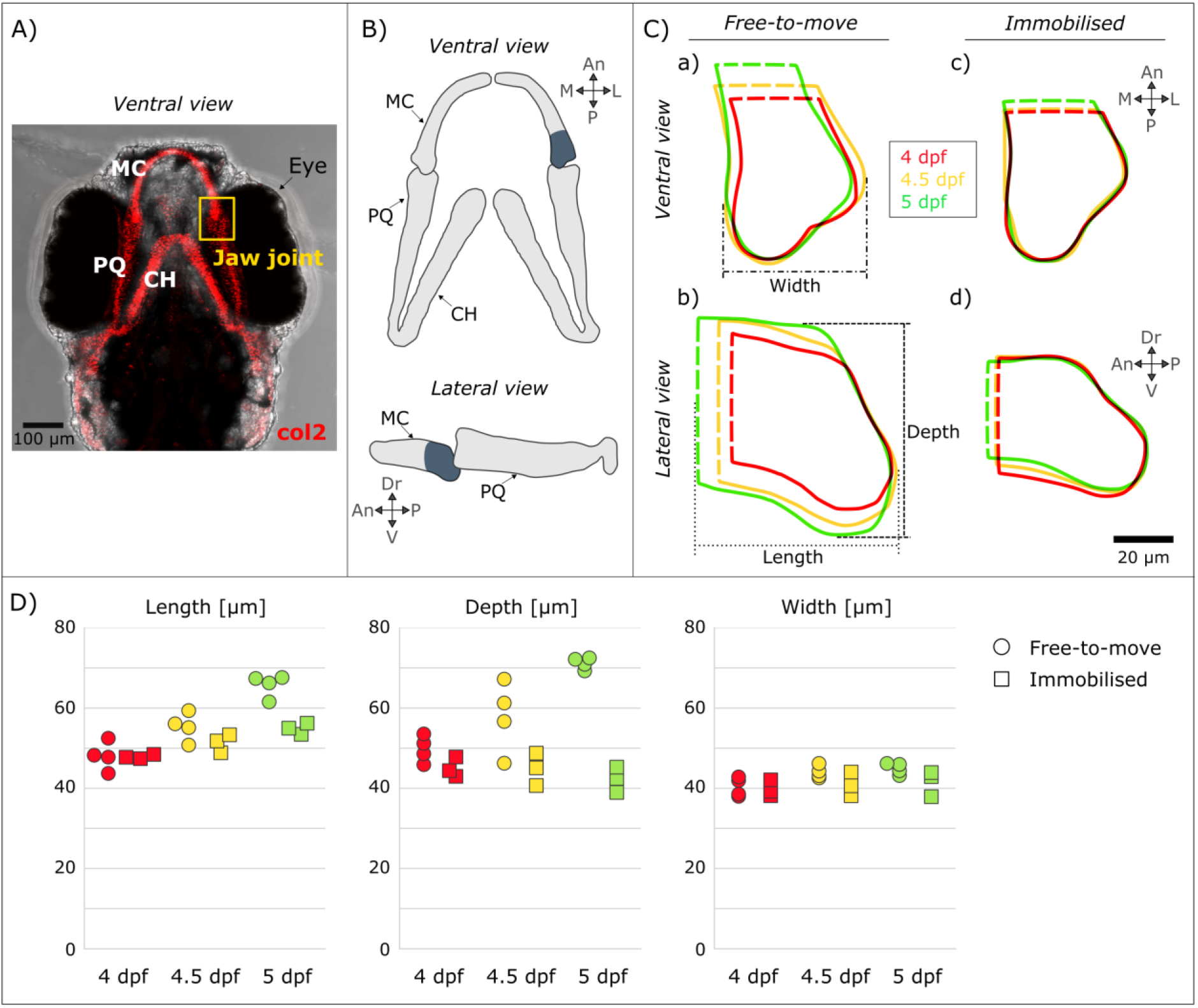
In the Meckel’s cartilage (MC) element, immobilisation affected growth in the depth more than growth in the width or length. (A) Brightfield ventral view of a 7 dpf zebrafish head expressing Tg(Col2a1aBAC:mCherry) cartilage marker showing the location of the jaw joint (yellow box). (B) Sketches of the jaw in the ventral and lateral planes illustrating the anterior Meckel’s cartilage (MC) element. (C) Shape outlines of average MC shape at 4, 4.5 and 5 dpf for free-to-move (a, b) and immobilised (c, d) larvae. (D) MC length, depth and width measurements taken on individual larvae from the free-to-move (n=4 per group) and immobilised larvae (n=3 per group) at 4, 4.5 and 5 dpf. An: Anterior, CH: Ceratohyal, Dr: Dorsal, L: Lateral, M: Medial, MC: Meckel’s cartilage, P: Posterior, PQ: Palatoquadrate, V: Ventral.

Growth rates were calculated from tracked cell-level data and visualised in the ventrodorsal, anteroposterior and mediolateral axes in free-to-move and immobilised larvae and over two time-windows: from 4 to 4.5 dpf and from 4.5 to 5 dpf. Ventrodorsal growth rates (along the depth) in immobilised larvae were significantly lower in both time-windows than in free-to-move larvae as shown in Fig 2A, B, E. Anteroposterior growth rates (along the length) were not different between free-to-move and immobilised larvae as shown in Fig 2A, C, F. Mediolateral growth rates were significantly lower in immobilised compared to free-to-move larvae from 4.5 to 5 dpf, as shown in Fig 2A, D, G, with the drop in the average mediolateral growth rate in the immobilised compared to the free-to-move group being less pronounced than for ventrodorsal growth rates over the same time window. To validate that the growth rates computed from tracked cell activities drive the observed shape changes, morphogenesis was simulated in free-to-move and immobilised larvae using finite element (FE) methods. The global patterns of free-to-move and immobilised jaw joint morphogenesis were correctly simulated using the growth rates obtained from cell-level data, including the observed depth increases in free-to-move controls but not in immobilised, and higher length increases in free-to-move controls than in immobilised, as shown in S1 Appendix. In conclusion, decreases in growth rates due to the elimination of jaw movements were most pronounced along the ventrodorsal axis, explaining the pronounced decreases in MC depth due to immobilisation. Therefore, immobilisation leads to growth rate alterations along specific anatomical axes meaning that growth anisotropy (the direction of tissue deformation) is altered when jaw movements are absent.

**Fig 2.**
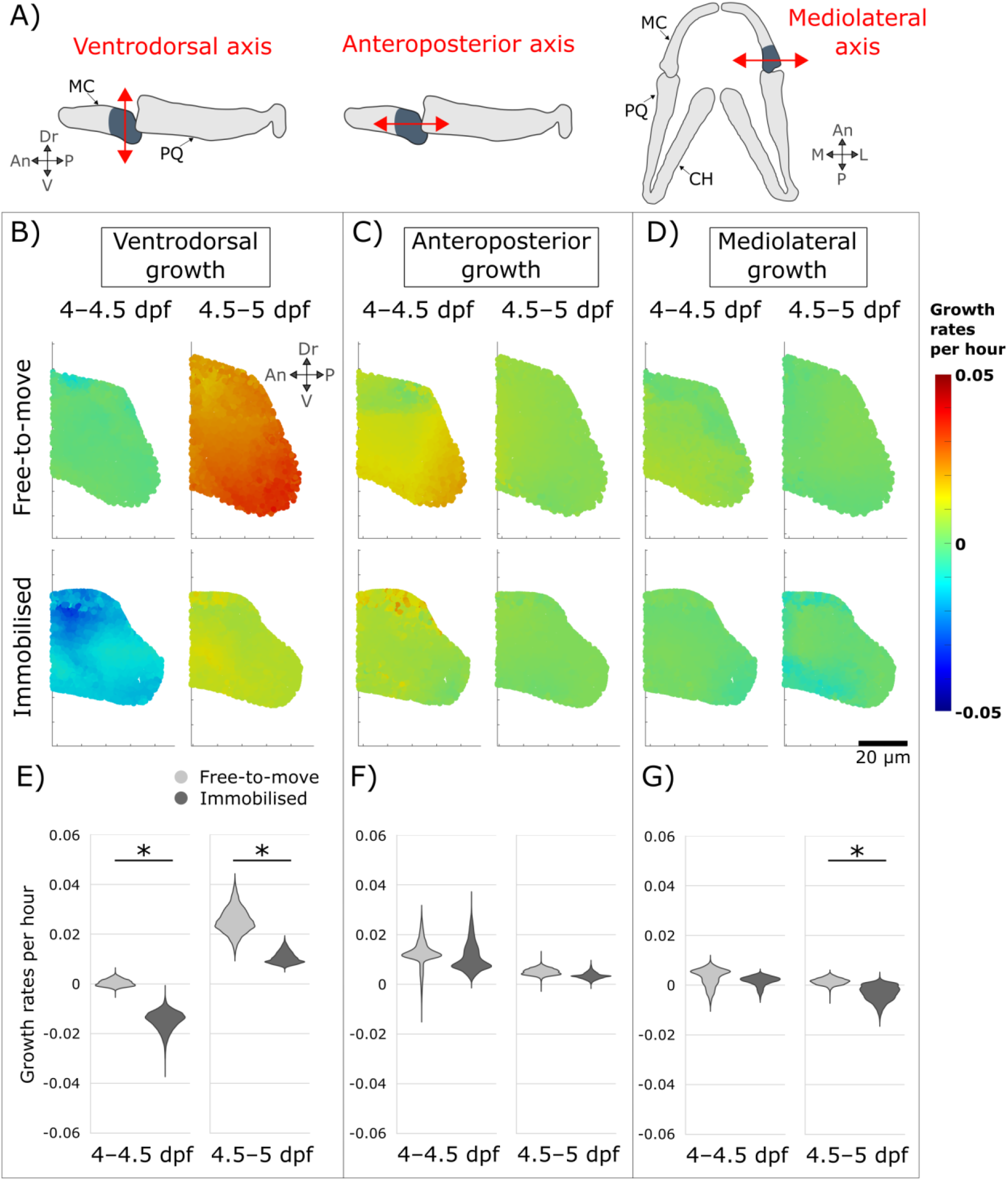
Immobilisation leads to altered growth rates, primarily along the ventrodorsal axis. (A) Illustration of the axes used for visualisation of growth anisotropy. (B, C, D) Ventrodorsal, anteroposterior and mediolateral MC growth rates from 4–4.5 and 4.5–5 dpf for both free-to-move and immobilised larvae. Results are displayed in one section in the mid-lateral plane. (E, F, G) Quantitative comparison of ventrodorsal, anteroposterior and mediolateral growth rates between free-to-move and immobilised groups. * indicates significant difference between free-to-move and immobilised means (p<0.05). An: Anterior, CH: Ceratohyal, Dr: Dorsal, L: Lateral, M: Medial, MC: Meckel’s cartilage, P: Posterior, PQ: Palatoquadrate, V: Ventral.

### Dynamic changes in load patterns-but not average values of compression-accurately simulate the mechanoregulation of jaw joint morphogenesis

In order to test hypotheses relating different types of mechanical stimuli to joint morphogenesis, we implemented simulations in which immobilised growth rates were applied to control joint shapes serving as the baseline biological contribution to growth. Then, a mechanobiological component of growth was combined to this biological baseline. Three different forms of mechanical stimuli arising from jaw movements were tested to determine which of them led to the most physiological pattern of growth; namely average hydrostatic stress, the dynamic switch from compression to tension over the loading cycle, and lastly the dynamic switch to and from compression without considering the influence of tension. Twelve-hour time intervals were simulated: from 4 to 4.5 dpf for which results are described herein, and 4.5 to 5 dpf whose results were consistent with the first time-interval and are therefore provided in S2 Appendix. The first stimulations used only the immobilised growth rates, representing the biological contribution only (**G**_**imm**_ in Figs 3A-i and S2A-i, red outlines in Figs 3A-iii and S2A-iii). For both time windows, the shapes grown under the biological contribution did not grow in depth, in contrast to when free-to-move growth rates were used (**G**_**free**_ in Figs 3A-ii and S2A-ii, green outlines in Fig 3A-iii and S2A-iii). Length increases were less pronounced using the immobilised growth rates (**G**_**imm**_, red outlines) than using the free-to-move growth rates (**G**_**free**_, green outlines) as shown in Figs 3A-iii and S2A-iii.

**Fig 3.**
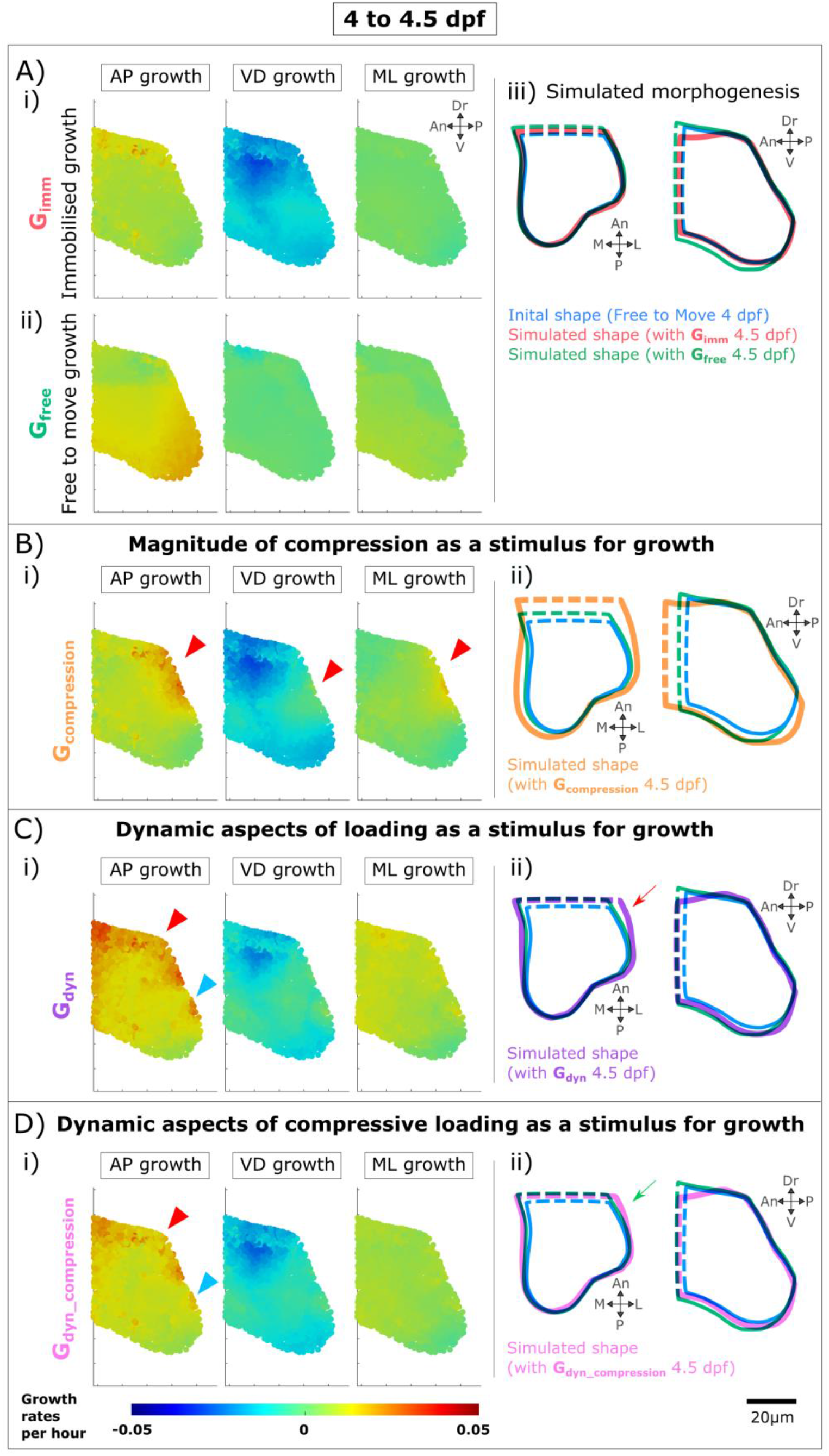
Biological and mechanobiological contributions to jaw joint morphogenesis from 4 to 4.5 dpf. A) Biological contributions to morphogenesis in the absence of movements lead to undergrowth of MC depth and length compared to free-to-move. A-i) Immobilised ventrodorsal (VD), anteroposterior (AP) & mediolateral (ML) growth rates applied to free-to-move 4 dpf shape. A-ii) Free-to-move growth rates. A-iii) Outlines of simulated morphogenesis with immobilised or free-to-move growth rates promoting growth. B) Average hydrostatic stress levels when used as the mechanoregulatory factor fail to simulate physiological jaw joint morphogenesis. B-i) Mechanobiological growth rates with compression promoting growth. Red arrowheads point to local areas of elevated growth rates which are not physiological. B-ii) Outlines of simulated morphogenesis. C) Dynamic patterns of mechanical stimuli (from compression to tension) lead to a more physiological growth pattern than average hydrostatic stress levels. C-i) Mechanobiological growth rates integrating the dynamic patterns of hydrostatic stress gradients. Red/blue arrowheads point local areas of elevated/reduced growth rates which are not physiological. C-ii) Outlines of simulated morphogenesis. Red arrow shows MC width overgrowth. D) Dynamic patterns of compressive mechanical stimuli lead to the most physiologically correct simulation of jaw joint morphogenesis. D-i) Mechanobiological growth rates integrating the dynamic patterns of compressive hydrostatic stress gradients. D-ii) Outlines of simulated morphogenesis. Green arrow shows the most physiological MC width as compared to previous simulations. An: Anterior, Dr: Dorsal, L: Lateral, M: Medial, P: Posterior, V: Ventral.

We first tested the hypothesis that joint mechanoregulated growth is proportional to the amount of compression experienced over joint motion and generated the mechanobiological growth map **G**_**compression**_. When simulating jaw opening and closing, average hydrostatic stresses were mostly spread in compression rather than in tension (Fig 4A), and a peak of compression was observed at the level of the jaw joint at both 4 and 4.5 dpf (Fig 4A, red arrowheads). For both time windows, the mechanobiological growth maps **G**_**compression**_ exhibited spots of locally increased growth rates at the level of the joint line which were not seen in the free-to-move growth maps **G**_**free**_ (red arrows in Figs 3B-i and S2B-i). From 4 to 4.5 dpf, mechanobiological simulations of joint morphogenesis using **G**_**compression**_ showed physiological MD depth growth compared to the shape grown under the biological contributions **G**_**imm**_, but overgrowth of both MC length and MC width compared to when the free-to-move growth rates **G**_**free**_ were used (Fig 3B-ii). From 4.5 to 5 dpf, there was physiological growth of MC length but no depth nor width increases using **G**_**compression**_ (Fig S2B-ii). Therefore, jaw joint mechanoregulated morphogenesis could not be accurately simulated using the hydrostatic stress levels averaged over joint motion as the mechanobiological stimulus.

**Fig 4.**
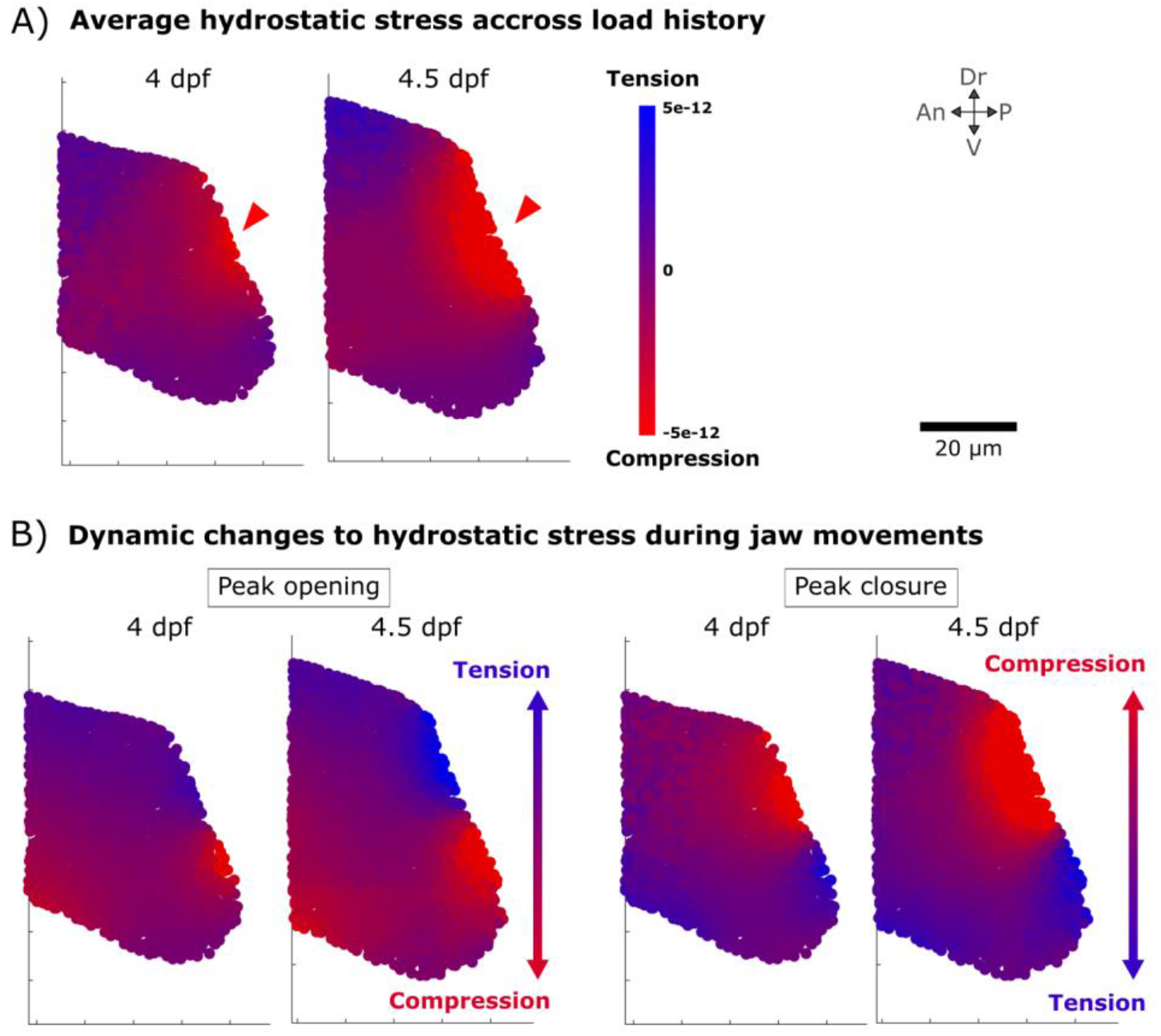
Compression and tension levels arising from jaw movements. A) Average hydrostatic stress distribution experienced over one cycle of mouth opening and closure at 4 and 4.5 dpf. Red arrowheads indicate a peak of compression at the level of the jaw joint. B) Dynamic changes to hydrostatic stresses from peak opening to peak closure showing a shift of hydrostatic stress gradients (from compression to tension) along the ventrodorsal axis. An: Anterior, Dr: Dorsal, P: Posterior, V: Ventral.

Since average hydrostatic stress levels as a stimulus for mechanoregulated growth failed to simulate physiological jaw joint morphogenesis, we next investigated the dynamic changes in hydrostatic stress patterns over jaw motion. At peak opening, the dorsal aspect of the rudiment experiences tension whereas the ventral aspect is in compression, creating a stress gradient from tension to compression along the ventrodorsal axis (Fig 4B). At peak closure, the dorsal aspect is in compression whereas the ventral aspect is in tension, creating a reversed gradient along the ventrodorsal axis compared to peak opening (Fig 4B). We hypothesised that the switch in the ventrodorsal hydrostatic stress gradient promotes growth along the rudiment’s depth and thus influences growth anisotropy. The time evolutions of hydrostatic stress gradients along the three anatomical axes were calculated and combined with **G**_**imm**_ to generate a new mechanobiological growth map called **G**_**dyn**_. From 4 to 4.5 dpf, simulating morphogenesis using the mechanobiological growth map **G**_**dyn**_ resulted in a shape which closely resembled the shape obtained using the free-to-move growth rates **G**_**free**_ (Fig 3C-ii, purple and green outlines). Increases in MC depth and length from 4–4.5dpf were almost the same between the **G**_**dyn**_ and **G**_**free**_ simulated shapes (Fig 3C-ii, purple and green outlines). MC width distant from the joint line was, however, bigger in the **G**_**dyn**_ simulated shape than in the shape obtained using **G**_**free**_ from 4–4.5 dpf (red arrow in Fig 3C-ii). From 4.5 to 5 dpf, mechanobiological simulations using **G**_**dyn**_ also resulted in a shape much like that obtained using the free-to-move growth rates, albeit slightly smaller (Fig S2C-ii, purple and green outlines). The mechanobiological growth maps **G**_**dyn**_ from 4–4.5 and from 4.5–5 dpf were similar to the free-to-move growth maps **G**_**free**_, all of which showed increased ventrodorsal (VD) growth rates compared to the immobilised growth maps **G**_**imm**_ (Figs 3C-i and S2C-i). A small number of differences between the free-to-move **G**_**free**_ and the mechanobiological **G**_**dyn**_ growth maps were observed. Anteroposterior (AP) growth rates were higher at the dorsal aspect in **G**_**dyn**_ in comparison to **G**_**free**_ where higher growth rates were observed at the ventral aspect for both time-windows (Figs 3C-i and S2C-i, red and blue arrowheads). From 4–4.5 dpf, **G**_**dyn**_ mediolateral (ML) growth rates were overall slightly higher compared to **G**_**free**_ explaining the MC width overgrowth (Fig 3C-i). Overall, from 4–4.5 and from 4.5–5 dpf, using the dynamic aspects of the hydrostatic stress fields arising from jaw movement as a stimulus for mechanoregulated growth enabled almost physiological jaw joint morphogenesis to be simulated.

Finally, we implemented simulations which included only the dynamic shift to and from compression without considering the influence of hydrostatic tensile stresses. For both time windows, simulated shapes when both tension and compression (**G**_**dyn**_) or only compression (**G**_**dyn_compression**_) were considered exhibited few differences as shown in Figs 3D-ii and S2D-ii. From 4-4.5 and 4.5–5 dpf, MC depth and length using **G**_**dyn_compression**_ were slightly smaller than when **G**_**dyn**_ was used but still alike that of the shape when the free-to-move growth rates **G**_**free**_ were used (Figs 3D-ii and S2D-ii, pink and green outlines). From 4–4.5 dpf, MC width increases were more physiological when only compressive stresses were considered (**G**_**dyn_compression**_) compared to when the influence of hydrostatic tensile stresses was considered (**G**_**dyn**_), (green arrow in Fig 3D-ii). Although the differences between both simulation types (**G**_**dyn**_ and **G**_**dyn_compression**_) are subtle, simulations using **G**_**dyn_compression**_ predicted shapes which resembled the most the shapes when free-to-move growth rates **G**_**free**_ were used. Therefore, the application of cyclic compressive loads is likely to be a major stimulus for mechanoregulated growth in the zebrafish jaw joint, while tension is probably not a key contributor to jaw joint morphogenesis.

## Discussion

In this research, growth patterns of zebrafish jaw joint morphogenesis were analysed and simulated in the presence or absence of movement. Growth when jaw movements were absent was most compromised along the ventrodorsal axis leading to pronounced decreases in MC depth in immobilised larvae compared to controls. Integrating cell-level data into mechanobiological models of jaw joint morphogenesis revealed that the dynamic patterns of mechanical stimuli arising from movements are more likely to stimulate mechanoregulated joint growth compared to the magnitude of loading alone. We showed that the application of cyclic compression, rather than the cyclical switch from compression to tension, is likely to be the key contributor to jaw joint morphogenesis.

We demonstrated for the first time that mechanical stimuli arising from fetal movements influence growth anisotropy in the developing joint. Chondrocyte orientation and intercalation have been shown to be affected when muscle contractions are absent in both fish [7, 15] and mice [6], and we propose that the effects on organ-level growth anisotropy we report could stem from these cell level changes. In support of this theory, computational models of limb bud elongation have demonstrated that anisotropic tissue deformation strongly influences the shape of the organ during chick [46, 49] and mouse [50, 51] hindlimb development, and that this anisotropy is correlated with patterns in cell orientations and with a bias in the orientation of cell divisions [51].

Previous mechanobiological models of joint morphogenesis have used a range of stimuli to promote growth and shape change, including average and peak hydrostatic stress [34-36, 38-41], peak octahedral shear stress [34, 35, 41], interstitial fluid pressure resulting from static or dynamic loading [37], or a combination of these. In the current research, when average hydrostatic stress distributions were used as promoters of mechanoregulated growth, morphogenesis of the zebrafish jaw joint was not accurately predicted. Rather, we found that the dynamic changes in the patterns of mechanical stimuli, and especially the cyclical application of compression, are the most likely stimuli influencing morphogenesis by altering growth anisotropy. This indicates that joint mechanoregulated growth is unlikely to be determined by solely the magnitude of mechanical stimuli experienced over motion. The importance of the dynamic nature of loading concurs with *in vitro* experimental data in which static loading downregulates chondrogenesis whereas dynamic loading upregulates it [21, 30, 31]. In the embryo, static loading—through rigid paralysis where the muscles are in continuous tetanus—disrupts joint morphology in larval zebrafish jaws and embryonic chick limbs [12, 15]. We propose that the dynamic nature of loading affects growth anisotropy and in turn morphogenesis, while acknowledging that the rates of growth are possibly influenced by the magnitude of mechanical stimuli.

A strength of this research is the direct incorporation of tracked cell-level data in the mechanobiological models when previous mechanobiological simulations of joint growth used extrapolated cell data and hypothesised how they impact growth rates. Previous computational simulations, including those from our group, assumed that the biological contributions to joint growth were proportional to chondrocyte density [36, 38, 52]. In this research, zebrafish jaw joint growth at the macro-scale was directly quantified from tissue geometry changes at the cellular level. This enabled precise and specific quantification of the cell-level growth and therefore less uncertainty when testing different hypotheses regarding to how mechanical stimuli influence growth.

There are some limitations to the current work. The zebrafish jaw joint has many similarities with mammalian synovial joints [53], but cavitation occurs later in development relative to the main events of morphogenesis in other animals including mice and humans [53]. However, this research investigates a critical time of joint morphogenesis, right after movements are established, and the advantages of the zebrafish (especially the transparency of the tissues enabling live cell tracking) outweigh its disadvantages. Another limitation of the zebrafish larval model when extrapolating to mouse or human is that the jaw joint has a very small number of cells and relatively low quantity of matrix in the tissue [48], and it is possible that individual cell behaviours have a greater impact on tissue shape than in organisms with more cells and proportionally more matrix. Therefore, our conclusions could be slightly altered in bigger animal models such as mammals. Another limitation is the use of linear elastic material properties when modelling zebrafish cartilage. Nano-indentation measurements showed that zebrafish cartilage rather exhibits viscoelastic properties. However, when jaw movement simulations were run using either material properties, inconsequential differences were observed.

In conclusion, in the absence of movement the directionality of growth in the joint is disturbed which affects joint morphogenesis. The magnitude of loading alone is not sufficient to explain the morphological changes observed at the organ-level during joint morphogenesis. Rather, changes in growth anisotropy are likely triggered by the dynamic changes in the mechanical stimuli experienced when cyclic compression is applied to the joint elements over joint motion. Overall, this research offers avenues for improvement in simulations of joint development and potentially other organs. It provides fundamental advance in our understanding of mechanoregulation in the developing joint and increases our understanding of the origins of conditions such as hip dysplasia and arthrogryposis.

## Materials and Methods

### Zebrafish husbandry, lines and anaesthetisation

Fish were maintained as described previously [54, 55]. All experiments were approved by the local ethics committee (Bristol AWERB) and performed under a UK Home Office Project Licence (PP4700996). Transgenic lines *Tg(col2a1aBAC:mCherry)* [56] and *Tg(−4*.*9sox10:eGFP)* [57] provide expression of fluorescent reporters for the immature chondrocytes in the interzone (*sox10-positive* and *col2-negative*) and the mature chondrocytes (positive for both *sox10* and *col2*). To study immobilised growth, wild type larvae were anaesthetised in 0.1 mg.ml^-1^ tricaine methanesulphonate (MS222) in Danieau’s buffer from 3 dpf prior to the start of recorded jaw movements [58]. The solution was refreshed twice daily until 5 dpf. Larvae were mounted in low melting point agarose (N.B. free to move larvae were briefly immobilised for image acquisition), and imaged on a Leica sp8 confocal.

### Characterising growth from cell-level data

Growth maps were calculated for 12-hour interval time windows (4–4.5 and 4.5–5 dpf) for free-to-move and immobilised specimens following the methodology previously published [48]. In free-to-move larvae consistent jaw movements are visible by 4 dpf [19]. Confocal image stacks centred on the jaw joint line were obtained at 4, 4.5 and 5 dpf for double transgenic *Tg(col2a1aBAC:mCherry; - 4*.*9sox10:eGFP)* free-to-move and anaesthetised larvae. In immobilised specimens, cells in the posterior palatoquadrate joint element could not be reliably segmented and tracked due to a weaker expression of fluorescent markers. Growth analyses were therefore performed solely on the anterior Meckel’s cartilage (MC) joint element. Cells were segmented in Fiji and the 3D cell centroid’s coordinates in the MC joint element were extracted at each time point [59, 60]. MC joint cells were manually tracked between images from two consecutive timepoints using manual labelling in MATLAB (R2018a, The MathWorks, Inc., Natick, Massachusetts, United States). The position of cell centroids with respect to each other over time was used to calculate the local rate of deformation in cubic regions of interests (ROIs) using the “statistical velocity gradient” equation from [61]. The cubic ROIs were of size length 15 µm and mapped the MC joint element. After calculations, the local growth in each ROI is represented by an ellipsoid whose axes represent the three directions for growth and their radius the rate of growth along these directions. Growth maps displaying the local deformation rates and directions were generated from the local growth ellipsoids for each time window. Interpolation between ROIs centres was performed in Abaqus CAE (Dassault Systemes, 2019) by importing the growth maps as analytical mapped fields (refer to “Simulating zebrafish jaw joint morphogenesis” section). In this study, ventrodorsal (VD)/anteroposterior (AP)/mediolateral (ML) growth is defined as the growth rates of the growth ellipsoid axis whose angle from the anatomical VD/AP/ML axis is the smallest amongst the three ellipsoid axes. All growth maps were analysed following this terminology. The growth ellipsoid axes are displayed in S3 Appendix, along with the angles between the growth ellipsoid axes and the anatomical VD/AP/ML axes. The number of samples analysed per time window are listed in Table 1. The p-value for growth rates mean differences along each direction for growth between free-to-move and immobilised groups for each time window were obtained by running Shapiro-wilk test of normally followed by Mann-Whitney U-test with Bonferroni adjustments for multiple comparisons.

**Table 1.** Sample number per time window for growth rates analyses in the anterior joint element of free-to-move and immobilised larvae.

|  | 4–4.5 dpf | 4.5–5 dpf | 5–5.5 dpf |
| --- | --- | --- | --- |
| Free-to-move | 7 | 7 | 7 |
| Immobilised | 8 | 6 | 7 |

### Average shape generation

Average shapes were generated at 4, 4.5 and 5 dpf for free to move and immobilised larvae following the methodology previously described [48]. Confocal image stacks of four to five larval zebrafish jaws (encapsulating the Meckel’s cartilage, the palatoquadrate and the ceratohyal) from the transgenic line *Tg(col2a1aBAC:mCherry)* were taken with a Leica SP8 confocal microscope at each time point in the ventral plane. A 3D Gaussian grey filter was applied to the image stacks in Fiji. Image stacks were imported in Mimics (Materialise NV, Leuven, Belgium) to be segmented. Only half-jaws (separated at the level of the midsagittal plane) were segmented. Each segmented half-jaw was divided into slices in the transversal plane. For each slice, an average shape outline was generated in MATLAB from the shape vertices of each segmented half-jaw. Averaged shape outlines were saved as image stacks and imported into Mimics where the resultant average half-jaw shape was generated. In the figures, the outlines of the average MC joint element were consistently cropped based on measured increases overtime of the distances between the tracked cells and the joint line.

### Material properties characterisation using nano-indentation

Jaw cartilage material properties in hypertrophic and immature regions were measured in wild type free-to-move and immobilised specimens at 4 and 5 dpf using nano-indentation. The indentation methodology was previously described [62]. Whole larvae were fixed in 4% PFA and stored in 100% MeOH. Prior to nano-indentation, samples were rehydrated to 1 x PBS before being stored in 30% sucrose in PBS. The samples were then submerged in a 1:1 mix of 30% sucrose and optimum cutting temperature (OCT) at room temperature until the samples sunk to the bottom of an Eppendorf tube. Samples were then embedded in fresh 30% sucrose and OCT mix and flash-frozen on dry-ice. Embedded samples were sectioned sagittally at a thickness of 10 µm using an NX70 Cryostat (CryostarTM, ThermoFisher, France). Nano-indentation was performed on sections featuring the Meckel’s cartilage (hypertrophic cartilage), the ceratohyal (hypertrophic cartilage) and/or the jaw joint (immature cartilage) using a Chiaro nanoindenter (Optics11 Life, The Netherlands) as shown in Fig 5A. All measurements were taken in PBS at room temperature. A spherical nano-indentation probe with an 8 µm radius and stiffness of 0.49 N/m was used. Indentation was performed to a depth of 1 µm with velocity of 1 µm/s, and the tip held at a constant depth for 10s (Fig 5B). Nano-indentation was performed across all sections containing relevant cartilage regions, with one measurement collected per region of interest in each section. This was performed for six larvae in each group except in the immobilised 4 dpf group where seven larvae were used. Young’s moduli were estimated using the Hertzian contact model, assuming a Poisson’s ratio of 0.3 (value which was previously used for AFM testing of the larval zebrafish jaw cartilage [62]). The resulting Young’s Moduli were averaged for each region across multiple sections of each fish. The Shapiro-Wilks test for normality was performed on each group. To test for significant differences between the hypertrophic and immature regions within each age and larva type group, paired t-tests were performed in groups which were normally distributed, and Wilcoxon signed-rank tests were used in groups which were not normally distributed. No difference was observed between hypertrophic and immature cartilage material properties as shown in S4 Appendix. Young’s moduli obtained from measurements taken in the immature regions are shown in Fig 5C.

**Fig 5.**
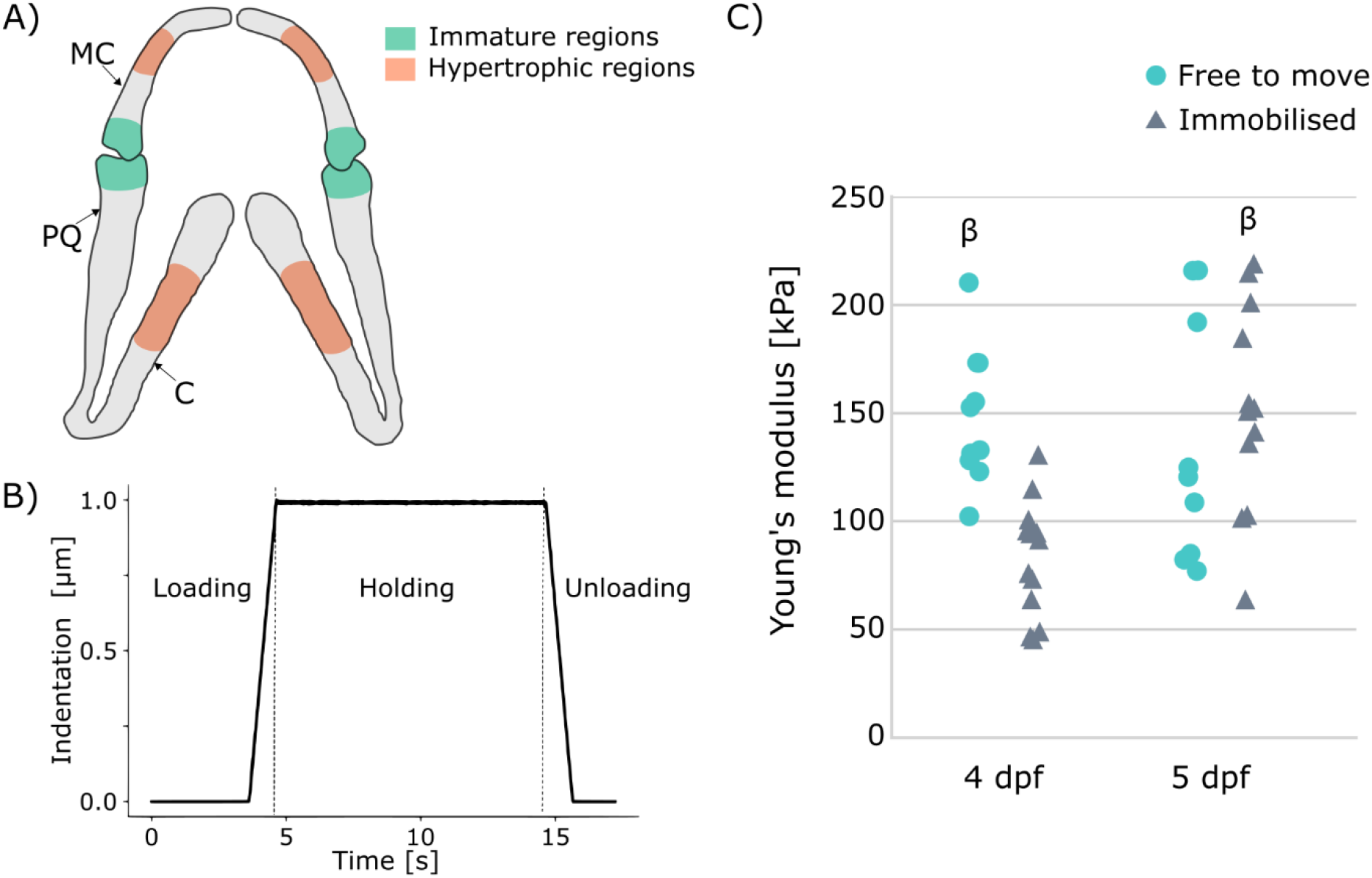
Indentation testing. A) Location of measurements in immature (green) and hypertrophic cartilaginous regions (orange) of larval lower jaw. MC: Meckel’s cartilage, PQ: palatoquadrate, C: ceratohyal. B) Indentation profile consisting of a loading phase during which a depth of 1 µm was reached, a holding phase of 10 s and an unloading phase. C) Young’s moduli of free-to-move and immobilised larvae at 4 and 5 dpf obtained from nano-indentation measurements taken in immature regions. β indicates significant difference (p<0.05) with 4 dpf immobilised group.

### Simulating zebrafish jaw joint morphogenesis

Morphogenesis in free-to-move and immobilised zebrafish jaws was simulated for each 12-hour interval time window (4–4.5 and 4.5–5 dpf) following the methodology previously described [48]. A non-manifold assembly combining the average half-jaw and the interzone (added as a volume filling the gap between the two joint elements using Boolean operations) was generated in Mimics and meshed in 3-matic (Materialise NV, Leuven, Belgium) with ten node tetrahedral elements. In Abaqus CAE, a finite element (FE) model for each twelve-hour time window was created. Cartilaginous regions were assigned homogeneous isotropic elastic material properties with Poisson’s ratio 0.3 and Young’s Modulus (YM) based on the nanoindentation measurements displayed in Table 2. Simulation tests were also performed using viscoelastic rather than linear elastic material properties (the tissue exhibiting viscoelastic behaviours) and resulted in inconsequential differences (S5 Appendix). No difference was observed between hypertrophic and immature cartilage material properties (S4 Appendix), therefore all cartilaginous elements were assigned the same material properties with no distinction between regions. The interzone was assigned isotropic elastic material properties with Poisson’s ratio 0.3 and YM set at 0.025% of the cartilaginous YM [62] and physiological boundary conditions were applied [48]. For each 12-hour period, growth strains derived from the growth maps were imported into Abaqus CAE as three distinct analytical mapped fields (one for each axis of the growth ellipsoids) and applied to the model. The Abaqus user subroutine UEXPAN was used to apply spatially varying expansion based on the strain fields to provide a prediction of growth and shape for each time-window. Outlines of the simulated shapes were obtained for the anterior MC joint element and cropped based on Abaqus results.

**Table 2.** Zebrafish jaw cartilage Young’s Moduli [kPa] in free-to-move and immobilised larvae based on nanoindentation measurements used in FE models.

|  | 4–4.5 dpf | 4.5–5 dpf | 5–5.5 dpf |
| --- | --- | --- | --- |
| Free-to-move | 142.01 | 142.01 | 142.01 |
| Immobilised | 82.91 | 117.44 | 151.96 |

### Simulating zebrafish jaw movements

Jaw movement simulations were performed on free-to-move 4 and 4.5 dpf FE models in Abaqus CAE. Muscle contractions engaged during opening/closure as shown in Fig 6 were applied to the models [63]. Muscle attachment points and directions were estimated from confocal scans of double transgenic *Tg(col2a1aBAC:mCherry;smyhc1:EGFP)* larvae (Fig 6). Muscle forces enabling physiological jaw displacement (jaw opening of 37.2 µm based on the average jaw displacement for 5 dpf larvae [19]) were used and are listed in Table 3. Jaw closure and opening were simulated in subsequent steps with each step decomposed into five increments. Hydrostatic stress and strain fields were extracted into MATLAB for each time increment from peak closure to peak opening.

**Table 3.** Muscle forces [nN] in the lower jaw enabling physiological movement in FE simulations.

|  | 4 dpf | 4.5 dpf |
| --- | --- | --- |
| adductor mandibularis (am) | 2.84 | 4.35 |
| intermandibularis anterior (ima) | 1.25 | 2.90 |
| intermandibularis posterior (imp) | 1.50 | 3.47 |
| interhyal (ih) | 1.50 | 3.47 |

**Fig 6.**
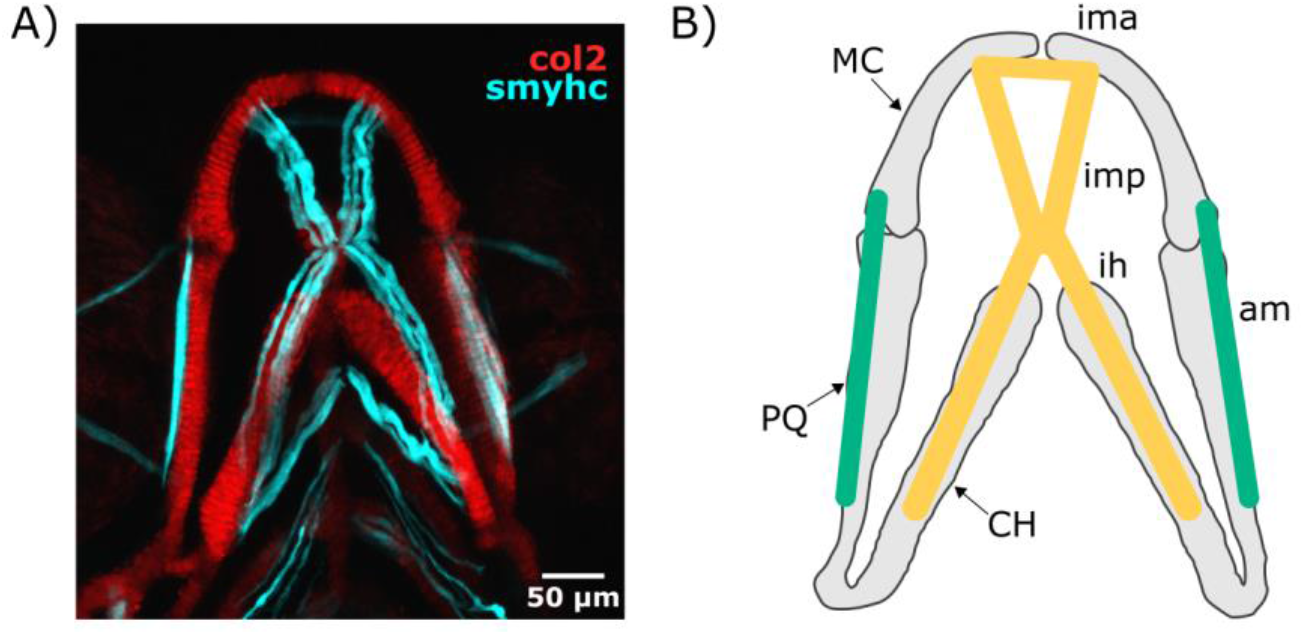
Lower jaw muscles. A) Maximum projection of ventral confocal image stacks expressing Col2a1aBAC:mcherry (red) and smyhc1:EGFP (cyan) of a 4 dpf larva. B) Scheme of the muscles engaged during lower jaw opening (yellow) and closure (green) in the ventral plane. CH: ceratohyal, MC: Meckel’s cartilage, PQ: palatoquadrate, am: adductor mandibularis, ih: interhyal, ima: intermandibularis anterior, imp: intermandibularis posterior.

### Investigating zebrafish jaw joint mechanoregulation

Loading fields were extracted from jaw movement simulations at 4 and 4.5 dpf and integrated into 4–4.5 and 4.5–5 dpf growth simulations respectively. The baseline levels of biological contributions to growth were assumed to be the growth rates calculated from immobilised larvae but applied to the free-to-move shapes (called **G**_**imm**_). Loading fields were used to alter **G**_**imm**_ and obtain new mechanobiological growth maps which were applied in growth simulations. Two different methods to calculate the mechanobiological growth maps were implemented to test different hypotheses. Since hydrostatic stress and strain patterns were similar (S6 Appendix), only the hydrostatic stress fields were used in the calculations for simplicity and uniformity with previous studies [34, 40].

First, we tested the hypothesis that the average hydrostatic stresses across loading history direct jaw joint morphogenesis with compression promoting growth. The average hydrostatic stress field across loading history ***S*** was calculated from the hydrostatic stress fields of all step increments in MATLAB:

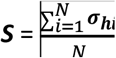

where ***σ***_***h***_ is the hydrostatic stress field and *N* the number of step increments from peak opening to peak closure. To test the hypothesis that compression promotes growth [24], a mechanobiological growth map **G**_**compression**_ was calculated based on the following equation (see Fig 7A and B):

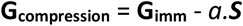

where *a* is a mechanoregulatory growth modulating variable which influences the impact of the average hydrostatic stress field ***S*** on the mechanobiological growth map. From 4 to 4.5 dpf, *a* was incrementally increased from 1e9 N^-1^s^-1^, a value with which it had minor influence on the mechanobiological growth map, to 4e9 N^-1^s^-1^ where joint overgrowth was obvious as shown in S7 Appendix. The modulating variable value which predicted physiological MC depth growth was chosen (*a* = 3e9 N^-1^s^-1^). The same value was used from 4.5 to 5 dpf.

**Fig 7.**
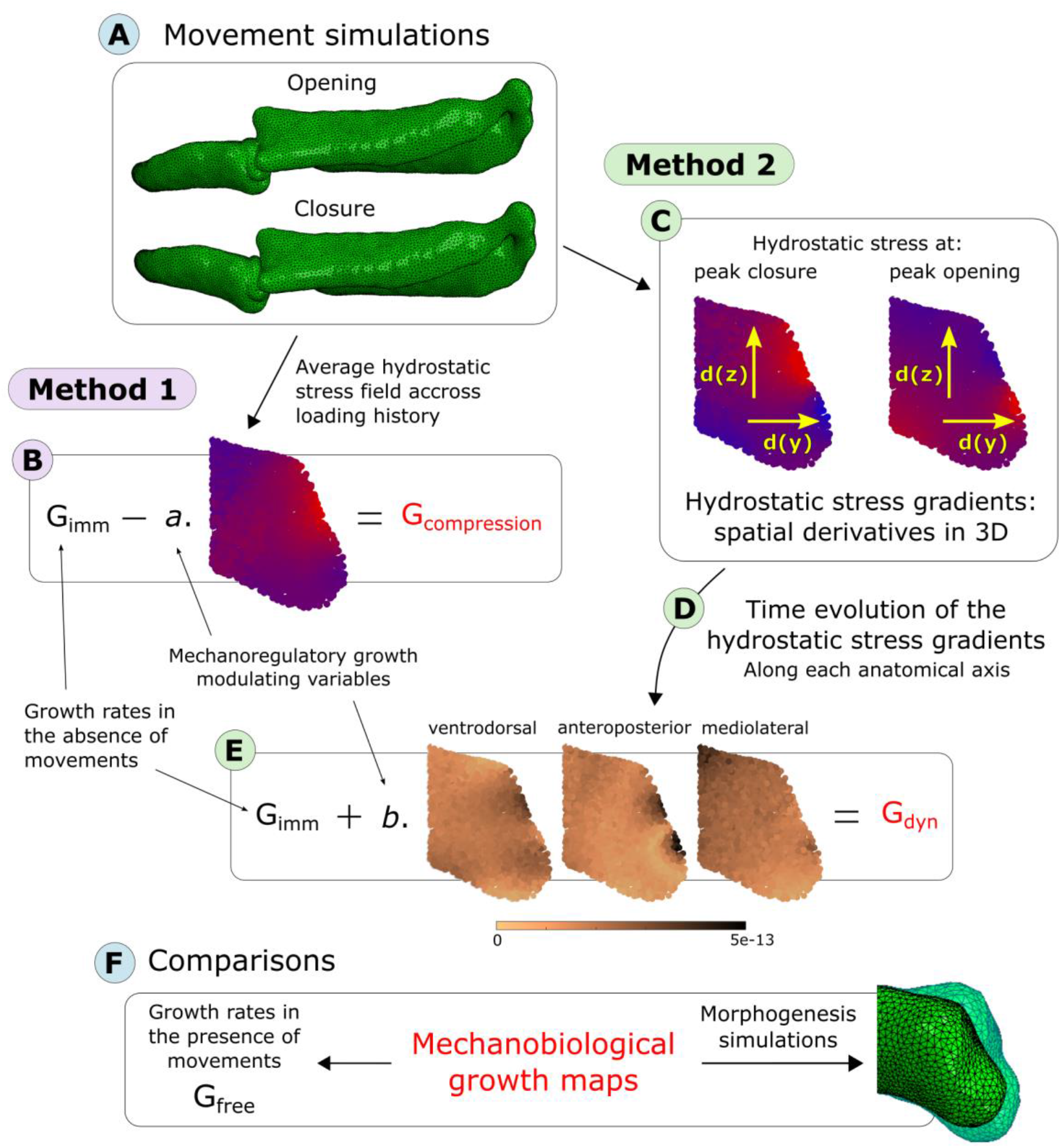
Process of integration of mechanical stimuli arising from movements into mechanobiological growth maps of zebrafish jaw joint morphogenesis. A) Jaw opening and closure were simulated. B) Method 1: Average hydrostatic stress field across the loading history was extracted and added to the growth rates in the absence of movements **G**_**imm**_ after being weighted by a mechanoregulatory growth modulating variable a. The newly obtained mechanobiological growth map was called **G**_**compression**._ C) Method 2: The time evolution of the hydrostatic stress gradients was used. Hydrostatic stress gradients at peak closure and peak opening were calculated along each anatomical axis. D) The time evolution of the hydrostatic stress gradients between peak opening to peak closure along each anatomical axis was calculated. E) and was added to **G**_**imm**_ after being weighted by a mechanoregulatory growth modulating variable b. The newly obtained mechanobiological growth map was called **G**_**dyn**_. F) The obtained mechanobiological growth maps **G**_**compression**_ and **G**_**dyn**_ were compared to the growth map in the presence of movements **G**_**free**_. Morphogenesis simulations were run using **G**_**compression**_ and **G**_**dyn**_ and compared to growth simulations using **G**_**free**_.

The mechanoregulatory growth modulating variable *a* encapsulates the number of jaw openings occurring over twelve hours (approximatively 57 thousand openings between 4 and 4.5 dpf [19]). The contribution of the average hydrostatic stress field to growth was isotropic: the same hydrostatic stress field and modulating variable was applied along all directions. The obtained mechanobiological growth maps **G**_**compression**_ was qualitatively compared to the growth map calculated from free-to-move larvae **G**_**free**_. Morphogenesis simulations were run in Abaqus CAE using **G**_**compression**_ based on the same methodology than explained in section “Simulating zebrafish jaw joint morphogenesis” and qualitatively compared to growth simulations using **G**_**free**_ (Fig 11F).

Next, we tested the hypothesis that <u>the dynamic changes in hydrostatic stress patterns over jaw motion direct jaw joint growth</u>. The hydrostatic stress gradients **∇ σh** at peak closure and peak opening were calculated along each anatomical axis in MATLAB, using the mathematical equations developed by [64] (Fig 7C). Along each anatomical axis, the time evolution of the pressure gradients between peak opening to peak closure was calculated: the absolute value of the difference between peak opening and peak closure was taken (Fig 7D). A new growth map **Gdyn** was calculated based on the following equation (see Fig 7E):

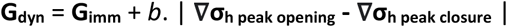

where *b* is a mechanoregulatory growth modulating variable which influences the impact of the hydrostatic stress field gradients on the mechanobiological growth map. From 4 to 4.5 dpf, *b* was incrementally increased from 1e10 m.N^-1^, a value with which it had minor influence on the mechanobiological growth map, to 1.5e11 m.N^-1^ where joint overgrowth was obvious as shown in S7 Appendix. The modulating variable value which predicted physiological MC depth growth was chosen (*b* = 5e10 m.N^-1^). The same value of *b* was used from 4.5 to 5 dpf. The number of jaw openings which occur over twelve hours is once again encapsulated in the modulating variable. The contribution of mechanical fields to growth was anisotropic: the same modulating variable value *b* was used along all anatomical axes, but the pressure gradients varied between axes. The newly obtained mechanobiological growth map **G**_**dyn**_ was qualitatively compared to the growth map calculated from free-to-move larvae **G**_**free**_. Morphogenesis simulations were run in Abaqus CAE using **G**_**dyn**_ and qualitatively compared to growth simulations using **G**_**free**_ (Fig 7F). To assess the <u>contribution of the application of cyclic compression to joint morphogenesis specifically</u>, we run more simulations where tensile hydrostatic stresses were left out. A mechanobiological growth map **G**_**dyn_compression**_ was calculated using the same methodology than for **G**_**dyn**_ except only the compressive hydrostatic stresses were considered. The same value for the mechanoregulatory growth modulating variable *b* was used (*b* = 5e10 m.N^-1^). Terminologies and methods for calculation of the mechanobiological growth maps are summarised in Table 4.

**Table 4:** Overview of growth maps terminologies and methods.

| Growth map | Description | Contribution of mechanical loads | Hypothesis tested | Mechanoregulatory growth modulating variable |
| --- | --- | --- | --- | --- |
| $G_{\text{free}}$ | Growth rates obtained from free-to-move cell-level data | Normal baseline | Growth simulations with tracked free to move cell data accurately predict free to move morphogenesis | - |
| $G_{\text{imm}}$ | Growth rates obtained from immobilised cell-level data | None | Growth simulations with tracked immobilised cell data do not accurately predict free to move morphogenesis | - |
| $G_{\text{compression}}$ | Mechanobiological growth maps | Average hydrostatic field across load history | Compression levels promote growth | $a = 3e9 \text{ N}^{-1}\text{s}^{-1}$ |
| $G_{\text{dyn}}$ | | Time evolution of the hydrostatic stress gradients | Dynamic load patterns promote growth | $b = 5e10 \text{ m.N}^{-1}$ |
| $G_{\text{dyn\_compression}}$ | | Time evolution of the compressive hydrostatic stress gradients | Dynamic patterns of compression promote growth | $b = 5e10 \text{ m.N}^{-1}$ |

All data underlying this article can be accessed on zenodo at https://doi.org/10.5281/zenodo.7586155. Confocal images, MATLAB scripts, Abaqus CAE models and simulated shapes are available.

## Acknowledgements

We thank James Monsen for providing the methodology and MATLAB script which was used for generating average shapes. We would like to thank Mat Green for zebrafish husbandry and the staff of the Wolfson Bioimaging centre Bristol for imaging support. We thank Dr. Labonte and his team for sharing nano-indentation equipment with us and Dr. Kaimaki for her valuable help during nano-indentation experiments.

## Supporting information captions

**S1 Appendix: Growth simulations from cell-level data**

**S2 Appendix: Biological and mechanobiological contributions to jaw joint morphogenesis from 4.5 to 5 dpf**

**S3 Appendix. Jaw joint growth orientations**

**S4 Appendix. Comparisons between the material properties of immature and hypertrophic regions**

**S5 Appendix: Comparison between linear elastic and viscoelastic material properties in jaw movement simulations**

**S6 Appendix. Comparison between hydrostatic strain and stress fields**

**S7 Appendix. Sensitivity analyses of mechanoregulatory growth modulating variables**

